# Short report: A judgment bias task reveals stress produces a negative affective state in cuttlefish

**DOI:** 10.1101/2024.04.15.589613

**Authors:** Sarah E. Giancola-Detmering, Robyn J. Crook

## Abstract

Judgment Bias Tasks (JBT) are used to assess emotional state and welfare of animals in zoos, farms, and laboratories, based on a subject’s interpretation of an ambiguous or intermediate cue. Animals in positive affective states are more likely to interpret the ambiguous cue positively, reflecting optimistic bias and animals with negative affect are more likely to interpret ambiguous cues pessimistically. Here, we developed a JBT assay for the stumpy-spined cuttlefish, *Sepia bandensis*, to determine whether cuttlefish exhibit negative affective states resulting from chronic and acute stress. Cuttlefish learned to associate food with a visual cue. Positive and neutral cues were presented twice daily until animals oriented to the positive cue and searched for food. After training, one treatment group was exposed to combined chronic and acute stress produced by 5 days of impoverished housing and once-daily simulated net capture. Our control group received no stress experience. In test trials performed after the stress experience, stressed animals had higher latencies to approach ambiguous cues, spent significantly less time in rooms with ambiguous cues once they entered, and were less likely to enter first into the ambiguous cue-paired room compared with controls. These behaviors suggest that stress induces pessimistic judgment bias in cuttlefish, the first indication of this capacity in cephalopods.

## Introduction

The use of cephalopods as model animals in biological research is growing rapidly. Their complex nervous systems, advanced cognitive abilities, and unique behaviors have made them ideal research models for neurobiology, behavior, and ecology. With the increase of popularity, the need for improved welfare and ways to assess stress levels is vital (Moltschaniwskyj et al., 2007).

One way of testing an animal’s welfare is to use cognitive bias (also known as judgment bias) to assess the animal’s emotional state (or affective state) based on how it views an ambiguous cue (Bethell, 2015). To use a judgment bias task, animals are trained to associate a positive cue with a positive reward and neutral/negative cue with no/negative reward. During the test phase, an ambiguous cue (in between the positive and neutral/negative cue) is presented. If the animal expects a reward in the presence of the ambiguous cue, it is in a more optimistic mindset or a positive affective state. However, if the animal treats the ambiguous cue as neutral, it’s in a more negative affective state or a pessimistic mindset (Harding et al. 2004).

JBTs have been used to assess cognitive capacity and welfare state in many vertebrate models such as rats, mice, dogs, zebrafish, horses and pigs (Rygula et al. 2015, Novak et al. 2015, Burman et al. 2011, Espigares et al. 2022, Freymond et al. 2014, Murphy et a. 2013). More recently, evidence is emerging that cognitive or judgment bias can be found in some invertebrates such as bumblebees and honeybees (Baracchi et al. 2017; Bateson et al. 2011). However, the use of JBT in invertebrates remains uncommon, and to our knowledge JBT has not been performed on a cephalopod. Chung et al performed a behavioral paradigm on *Sepia pharaonis* inspired by a JBT with hopes to find evidence of affective states. This was not a proper JBT since there was no training phase with a positive or neutral/negative cue. Instead, it relied on the cuttlefish’s prey choice through a discrimination task. Ultimately, this did not work and they recommended using alternative methods to assess affect in cuttlefish.

Evidence for affective states in cephalopods is accumulating. It has been up for debate whether cephalopods are sentient, therefore able to experience different affective states. One possible sign of a positive affective state is play behavior. Not only does play show signs of higher cognitive abilities, but it’s also a proxy for proper welfare and positive affect (Ahloy-Dallaire et al., 2018). When an animal can play, this means their necessary needs are being met and they are able to engage in this behavior, showing signs of positive affect (Boissy et al., 2007). This play behavior has been shown in other cephalopods such as *Octopus vulgaris* and *Octopus dofleini*, where they’ve been shown to push a floating plastic pill bottle at the top of the tank with their siphon and play-like behavior with a Lego and a plastic bottle (Mather & Anderson, 1999; Kuba et al., 2006; Kuba et al., 2003). Negative affect has also been observed. In a recent study, *Octopus bocki* showed conditioned place avoidance when injected with painful acetic acid, indicating not only pain experience but a negative affect (Crook, 2021).

Although previous studies of negative affects in cephalopods did not address stress, recent evidence suggests that non-painful, stressful experiences do cause long-lasting changes to physiology in cephalopods (Chancellor et al., 2023). Because little is known about optimal or stress-limiting housing, handling, and management of cephalopods in captivity, it is likely that at least some routine procedures are experienced as stressful. Little to no enrichment for an animal can also cause stress (Fox et al., 2006). For cuttlefish specifically, research shows that impoverished housing causes a slower growth rate and negatively affects their memory and cognition (Dickel et al., 2000). Similar work has also been shown that having an enriched environment improves their cryptic coloration (Poirier et al., 2005), and increases hunting success (Yasumuro & Ikeda, 2018).

Chronic and acute stressors have been shown to negatively affect vertebrate animals in captivity, causing cognitive and behavioral changes (Mendl, 1999). There is a wide range of behaviors that reveal stress in vertebrate animals such as, repeated movements (Ferdowsian et al., 2012), decrease in exploratory behavior (Vyas and Chattaji, 2004), excessive grooming (Komorowska & Wojciech, 2003), and changes in social and sexual behavior (Hemsworth et al., 1986). Chronic as well as acute stress can cause negative changes to learning and memory. Multiple studies have shown vertebrate animals exposed to stressful stimuli, enrichment removal, and unpredictable housing conditions, caused animals to respond negatively (or pessimistic) to an ambiguous cue during judgement bias trials (Harding et al., 2004; Bethell, 2015; Brydges & Hall, 2017), indicating both that these events are stressful and that JBT is an appropriate assay to capture animals’ experiences of them.

This study used the Go/No-Go JBT with visual cues. Cuttlefish were trained to approach paddles marked either with vertical or horizontal stripes for a food reward, and not approach a neutral paddle with no reward (Bethell, 2015). Given the large volume of studies using visual cues to direct cuttlefish behavior (Hvorecny et al., 2007; Barbosa et al., 2012; Scatà et al., 2016), we expected that this task would be readily performed by our animals. We used a combination of chronic exposure stress and acute handling stress to attempt to produce a transient negative affective state and assessed whether these treatments produced evidence for pessimistic judgement bias.

## Materials and Methods

### Animals

Stumpy-spined cuttlefish or dwarf cuttlefish *(Sepia bandensis, N=17)* were captive-bred and purchased as hatchlings from Marine Biological Laboratory Center for Cephalopod Culture (Massachusetts, USA). Cuttlefish were fed *ad libitum* on live mysid shrimp (*Mysidopsis bahia)* until about 6 weeks post hatching. After 6 weeks, they were fed 3 live grass shrimp (*Paeneus spp*.) per day. Cuttlefish were maintained in a recirculating seawater system (1600 L) held at 23.5-25.5 C and filtered via physical, chemical, and biological filtration. Cuttlefish were reared in floating tub enclosures (30 cm diameter and 8 cm deep) with 4-5 hatchlings/tub until animals were separated and housed individually for trials. Each housing tub contained a sand and pebble bed and various enrichments (plastic plants, coral rubble, shells, and PVC tubes, Figure 1A&B). Cuttlefish reared in these conditions reach sexual maturity at around 3.5 months and a maximum size of 70mm (mantle length). For both sexes, stumpy-spined cuttlefish have a life span of 7-10 months. Experiments were conducted between May 2023-March 2024.

**Figure 1.**
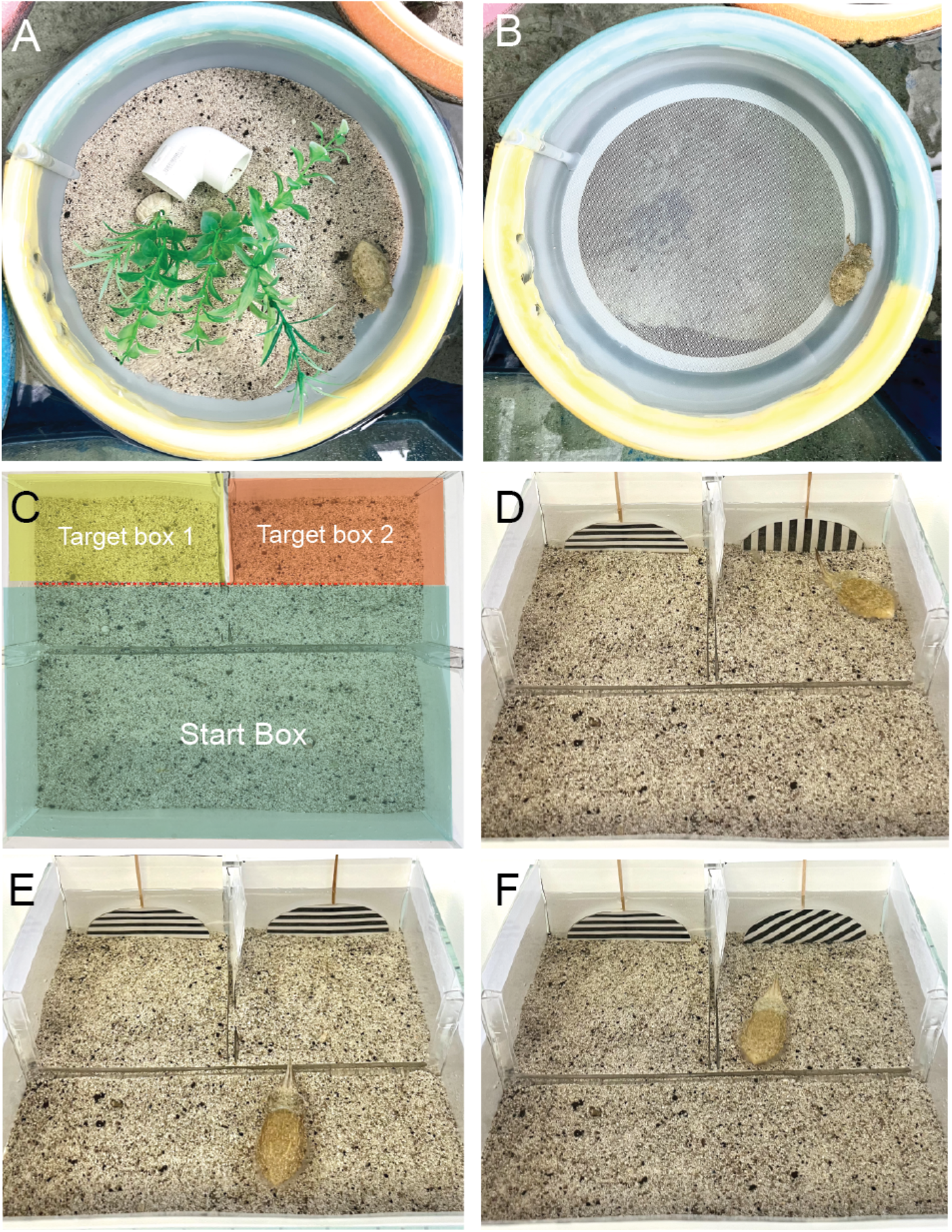
(a) Shows a standard, enriched housing setup which we used as our control condition. (B) Housing setup for treatment animal with impoverished housing. (C) Shows picture of y-maze set up with red lines and shading to show the two target rooms (or boxes) signaled by a cue on the back wall. When swimming forward, the eyes of the animal had to pass the red line to count as an entry. Dimensions of each room 7.5x10.3 cm. (D) Shows y-maze set up for alternating sides training trials (with frozen shrimp next to positive paddle) with positive and neutral paddle. (E) picture shows y-maze set up for trials for double neutral training trials with two neutral paddles. (F) Picture shows y-maze set up for trials for AMB trials with 45° diagonal line and a neutral paddle. For all trials, “correct” sides were assigned to the left or right box randomly, so animals learned the cue and not the location.

### Pre-Training

For two weeks cuttlefish were trained to start recognizing a cue (either a horizontal or vertical striped paddle) with a positive food reward. This was done by setting a paddle in the tub with stripes at exactly 180 degrees (horizontal) or 90 degrees (vertical) when food was given.

During these two weeks cuttlefish were also trained to eat frozen shrimp by attaching a frozen shrimp to a fishing wire and dangling it in front of the cuttlefish. If the cuttlefish did not eat the frozen shrimp then a day of feeding was skipped (never exceeding 2 days). All cuttlefish learned rapidly to take frozen shrimp, which were placed onto the enclosure floor. Cuttlefish moved on to the next step when they were showing orientation toward the visual cue and hunting behavior toward the frozen shrimp.

Hunting behavior was defined as the cuttlefish orienting its body towards the visual cue, swimming slowly towards it with a postural component go front raised arms (usually darkened) and waving side to side (Adamo et al., 2006, Hanlon and Messanger, 1988). In this study we term this suite of behaviors “hunting body pattern”.

### Training Trials

The experimental apparatus was a Laden Glass Air Aquarium 1.6-gallon rimless tank (10.6 x10.6 x 4.7). The tank was placed on a stage with two Ulanzi Ultra Bright LED Video Lights providing illumination of the arena from above and each side, with a Sony AX33 4k Handycam camera recording directly overhead. The tank was turned into a three-chamber Y-maze by adding a wall halfway, and a shorter divider forming two small target boxes (Fig 1C). There was a barrier placed in the middle of the aquarium that could be removed to start the trial. At the back of the two-chamber rooms, the positive cue and the neutral cue were placed against the rear wall, with the cues positioned at 180 and 90 degrees (Figure 1D).

The cuttlefish was placed in the start room with the middle barrier in place to obstruct the view of the cues. After two minutes of acclimation the middle barrier was removed and the cuttlefish could view cues in each room.

For the first 3 trials, full frozen shrimp were dropped in front of the positive cue to get the attention of the cuttlefish. Positive cue position was randomized to the left or right chamber, so the cuttlefish did not associate a side with a positive food reward. For the following trials, two frozen shrimp were placed at the back of the reinforced chamber right up against the cue, with the neutral chamber having nothing in the room except the cue paddle. Each trial ran for 10 minutes, or until the cuttlefish successfully ate the shrimp.

A minimum of 8 trials were conducted before the individual could move to the next phase. The criteria to move to the next phase were 1. Cuttlefish must successfully choose the correct side first in 2 of the last 3 trials and 2. Cuttlefish must only search the correct room in 2 of the last 3 trials. In other experiments there are many criteria to pass this training session such as having a 60-90% correct responding to the positive and neutral cues (Anderson et al., 2013; Keen et al., 2014; Rygula et al., 2015). Cuttlefish that did not reach the 2 criteria continued the training trials until reaching the criteria or reaching to 10 trials of training and were thus excluded. This process is similar to those seen in other studies (Starling et. al., 2014; Verbeek et al., 2014; Bethell and Koyama, 2015).

### Double Neutral Trials

To assess successful acquisition of the task, once criterion was reached we performed a single trial with both chambers containing the neutral cue and no food rewards offered (Figure 1E). These “Double Neutral” or DN trials were otherwise identical in procedure to training trials. We considered these trials as successful if the animals walked up to the room and either chose not to enter, or entered with increased latency compared with the most recent training trials. If cuttlefish did not meet at least one of these criteria, the cuttlefish went back to training, and must have re-met the training criteria over three successive trials, then they were retested on DN trials. All animals in the study passed the DN phase on either their first or second attempt. Before proceeding to the test trials, we gave a single reminder training that was identical to all other training trials.

### Stress experience

The treatment group was given impoverished housing (removing all enrichments (Figure 1 A, and leaving a bare tub, Fig. 1B) 6 days before the trials. 3 days before trials, the treatment group were chased and repeatedly briefly restrained by an experimenter with a small hand net for 3 minutes, twice per day. Cuttlefish showed several signs of being stressed during the 6-day period; during the impoverished housing period, the cuttlefish would sometimes take longer to eat their live shrimp, whereas unstressed cuttlefish generally swam right for the shrimp immediately after the shrimps were dropped into the tub. This suggests reduced food motivation, which is one sign of chronic stress. When the net would gently chase the cuttlefish, they also showed signs of acute stress by either sticking to the side of the tub with increased respiration or would jet away and ink. Stressed animals also showed changes to body patterns, almost always dark in coloration and dark around eye.

### Test Trials

Test trials took place within the same experimental apparatus set up as the training and double neutral trials. The two cues placed in the back of the two-chambered rooms were the existing neutral cue, and a new “ambiguous cue”, with black and white lines at 45° diagonal (i.e., exactly intermediate between the horizontal and vertical cues). No frozen shrimp were placed in the end boxes (Figure 1F). Cuttlefish were placed in the start room with the middle barrier in place. Cuttlefish were placed for two minutes for acclimation (within 1 minute the camera was turned on to start recording). After two minutes, the middle barrier was removed, and cuttlefish could see both cues in each room. Each test trial ran for 10 minutes, identical to other trials.

Animals were fed directly after every test trial to ensure that any lack of food search behavior was not due to a lack of food motivation - all animals ate readily once returned to their home tanks. We performed two test trials over consecutive days for each animal. A diagram timeline of the full experiment is shown in Fig 2.

**Figure 2.**
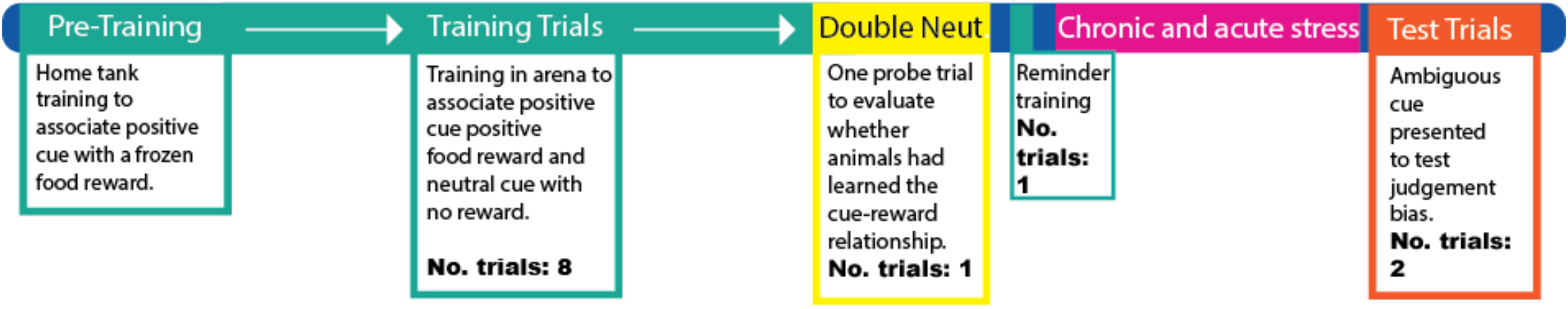
Timeline diagram showing the sequence of trial types and the number of each kind.

### Data Analysis and Statistical procedures

From recorded footage of each 10-minute trial, we recorded side first entered, latency to enter first room and correct (reinforced) room, latency for cuttlefish to choose ambiguous room (in test trials), duration of time spent in each room, and duration of time spent exhibiting hunting body pattern.

“Side first entered” was defined as the room the cuttlefish moved into first after the trial was started by removal of the horizontal barrier. Entry was counted when the animal’s eyes had crossed the “room line” (see figure 1c, for room delineation). We used unpaired (independent) t-tests to compare the animals in each group to the average. The critical alpha was set at 0.05, and all P-values were two-tailed.

For latency to enter the first room, time started once the middle divider was removed and stopped once cuttlefish eyes crossed the “room line.” For the training trials the last two trials were averaged. Only a single Double Neutral (DN) trial was performed, so the values shown are from a single trial per animal. Latency for the test trials was averaged between the first and second day of trials. We used unpaired (independent) t-tests to compare the animals in each group to the average. The critical alpha was set at 0.05, and all P-values were two-tailed.

Latency to enter the ambiguous room was calculated for test trials only. Time started once the divider was removed and stopped when the cuttlefish entered the ambiguous room (measured as the point at which the eyes crossed the room line). This included time spent in another room chosen first until they chose the ambiguous room. Latencies where the animal stayed in the start room were recorded at 600 seconds (10 minutes). Times were averaged across the two tests given on consecutive days. We used unpaired (independent) t-tests to compare the animals in each group to the average. The critical alpha was set at 0.05, and all P-values were two-tailed.

Time spent in a room while expressing hunting body pattern was also measured for each test trial. This meant once the animal chose a room (either ambiguous or neutral), crossed the “room line” and was expressing hunting body pattern, the time was recorded. If the hunting body pattern stopped, or they exited a room, then the timer stopped as well. Time in HBP for each room was averaged between both trial days. We used unpaired (independent) t-tests to compare the animals in each group to the average. The critical alpha was set at 0.05, and all P-values were two-tailed.

## Results

Latency to enter either target box was measured for training, DN and test trials. During the two final training trials the latency to enter the correct room was observed and averaged. There was no significant difference between the treatment group and control (figure 3A). During the double neutral trial there was also no pre-existing significant differences between the control and stressed animals. After 6 days of stress, latencies from the first and second day of test trials with the ambiguous cue were averaged, and the treatment group showed a significantly higher latency than the control group to choose a room to enter (Fig 3A, unpaired (independent) t-test, p=0.0370). We also measured the latency to enter the ambiguous room during the test trials (rather than either room), and show that stressed animals take significantly longer to enter the a room with an ambiguous cue compared with controls (Fig. 3B) unpaired (independent) t-test, p=0.0147). Averaged over both days of test trials, control cuttlefish spent significantly more time in the ambiguous room searching with HBP vs the stressed group (Fig 3C, unpaired (independent) t-test, p=0.0081).

**Figure 3.**
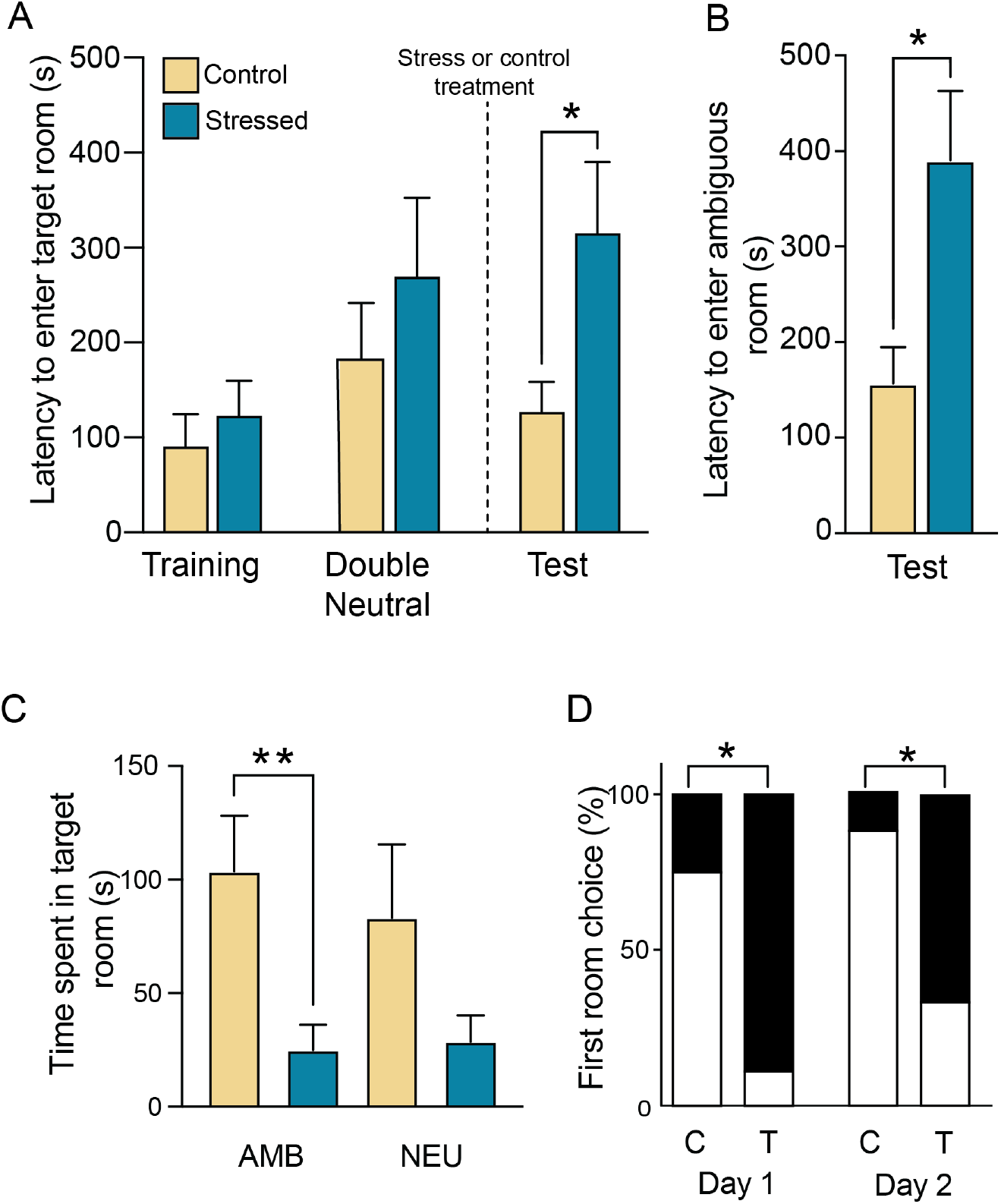
(A) Latency to enter either of the target rooms. For each animal, we averaged the final two successful training trials, there was a single double neutral trial, and then both test trials were averaged. Bars show group means and error bars are ± SEM. In test trials conducted after the stress or control treatment, animals in the stressed group showed increased latency to enter either target box (ind. sample t-test, p=0.0370). B. Latency to enter the room with the ambiguous cue, presented for the first time in test trials. Results from both test days are averaged for each animal (ind. sample t-test, p=0.0147). (C) Total time spent in each room with HBP (p= 0.0081). Stressed animals spent significantly less time searching for food in the room with the ambiguous cue. Although the also spent somewhat less time in the room with a neutral cue, this difference was not significant. Bars show group means and error bars are ± SEM. D. We compared the proportion of animals in each group that chose to enter the room with the ambiguous cue first. Data are shown for each test day separately. On both test days, a significantly greater percentage of control animals (C) chose the ambiguous room (white) to enter first. In contrast, the majority of treatment-group cuttlefish (T) chose a different room (either never left the start box or entered the neutral room first. Fisher’s Exact tests of C vs T, Day 1, p=0.015, Day 2, p=0.049)

For counts of choices of first room entered (binary outcomes) we analyzed the first and second day of testing separately. On the first day, Control cuttlefish were significantly more likely to enter the ambiguous room first, compared with treatment cuttlefish (Fig 3D, Fisher’s exact test, p=0.015). The second day of test trials show similar trends with control cuttlefish choosing ambiguous room first significantly more than the treatment, (Fig 3D, Fisher’s Exact test, p=0.049).

## Discussion

Here we demonstrate that cuttlefish show evidence of cognitive bias, the first time this ability has been tested in any cephalopod. The combination of acute and chronic stress likely represents common experiences of laboratory-housed cuttlefish; thus our data also show that cuttlefish can experience negative affective states as a result of sub-standard housing and handling approaches. In general, control group cuttlefish were more likely to choose to enter the room containing the ambiguous cue compared with treatment-group animals. They also had a significantly shorter latency to enter the ambiguous room during the test trials and spent more time in the room containing an ambiguous cue. Together, these data suggest control-group animals interpret the ambiguous cue in a more positive light compared with stressed animals, and therefore that control cuttlefish were experiencing a more positive or optimistic affective state. In contrast, the stressed cuttlefish show signs of treating the ambiguous cue more like the unrewarded neutral cue. Many animals in the stress group chose either the neutral room or stayed in the start room during test trials. This behavior was infrequent in the control cuttlefish during the test trials, and in any cuttlefish during the training trials. It is possible that stress, along with affecting judgment and emotional state, may also affect attention and memory recall (Mathews et al. 1995; Mineka and Sutton, 1992). We included a second test day to increase the chance of capturing reliable test data from each animal and results were consistent across both days, suggesting that transient effects of stress on memory were less likely to explain our results. Suppression of feeding motivation as a result of stress was also unlikely to explain our data, as all animals in the stressed group ate readily and immediately when returned to their home tanks and had been eating normally throughout the previous five days of the stress treatment.

We also considered that cuttlefish may not have learned the task completely by the time the test trials were conducted. We included a “Double Neutral” trial prior to beginning the stress or control treatments to evaluate how well animals had learned to associate approaching the reinforced cue with a food reward. Some cuttlefish during the double neutral trails would walk up and look at both cues but never choose a room. This suggests they had learned both cues were neutral, and as such would not have a food reward. This could be how the stressed cuttlefish viewed their options during the test trials, interpreting the ambiguous cue to be a neutral cue and choosing not to perform food search behaviors. Their increased latency to choose a room in the test trials also suggests lower motivation to investigate the room containing the ambiguous cue.

These data suggest acute and chronic stress have negative effects on cephalopods that are not only physiological (Chancellor et al., 2023) but also cognitive and affective. To our knowledge, this is the first indication of cognitive or judgement bias in cephalopods, adding to the small but growing body of literature suggesting that this capacity for complex processing is not exclusive to vertebrates (Crook, 2021; Baracchi et al., 2017; Kuba et al., 2006; Bateson, 2011). For laboratories housing cuttlefish, our data show that if enclosures and housing do not have sufficient enrichment, or if animals are exposed to acute stress during handling, this may affect animal welfare and potentially also results of experiments. While this is problematic for laboratory-housed cuttlefish, it also suggests that the many cuttlefish kept in zoos and aquariums may experience poor welfare if their housing and husbandry contains similar stressors. We suggest that JBT is useful and novel tool for assessing welfare of captive cephalopods, and may be valuable for assessing the effects of refinements and improvements to their care in laboratory and educational settings.

## Acknowledgements

This study was funded by NSF IOS CAREER 2047331 to RJC. We thank all members of the Crook Laboratory for assistance with experimental procedures, animal care and husbandry.

